# Conformational ordering of intrinsically disordered peptides for targeting translation initiation

**DOI:** 10.1101/2020.08.20.230268

**Authors:** Christopher J Brown, Chandra S Verma, David P Lane, Dilraj Lama

**Affiliations:** p53 Laboratory, A*STAR (Agency for Science, Technology and Research), 8A Biomedical Grove, #06-04/05, Neuros/Immunos, Singapore 138648; Bioinformatics Institute, A*STAR (Agency for Science, Technology and Research), 30 Biopolis Street, #07-01 Matrix, Singapore 138671; Department of Biological Sciences, National University of Singapore, 14 Science Drive 4, Singapore 117543; School of Biological Sciences, Nanyang Technological University, 50 Nanyang Drive, Singapore 637551; Department of Microbiology, Tumor and Cell Biology, Karolinska Institutet, Biomedicum Quarter 7B-C Solnavägen 9, 17165 Solna, Sweden

## Abstract

Intrinsically disordered regions (IDRs) in proteins can regulate their activity by facilitating protein-protein interactions (PPIs) as exemplified in the recruitment of the eukaryotic translation initiation factor 4E (eIF4E) protein by the protein eIF4G. Deregulation of this PPI module is central to a broad spectrum of cancer related malignancies and its targeted inhibition through bioactive peptides is a promising strategy for therapeutic intervention. We have employed a structure-guided approach to rationally develop peptide derivatives from the intrinsically disordered eIF4G scaffold by incorporating non-natural amino acids that facilitates disorder-to-order transition. The conformational heterogeneity of these peptides and the degree of structural reorganization required to adopt the optimum mode of interaction with eIF4E underscores their differential binding affinities. The presence of a pre-structured local helical element in the ensemble of structures was instrumental in the efficient docking of the peptides on to the protein surface. These insights were exploited to further design features into the peptide to propagate bound-state conformations in solution which resulted in the generation of a potent eIF4E binder. The study illustrates the molecular basis of eIF4E recognition by a disordered epitope from eIF4G and its modulation to generate peptides that can potentially attenuate translation initiation in oncology.

## Introduction

A significant population of eukaryotic proteins are found to contain continuous stretches of amino acids (> 30) that lack a well-defined tertiary structure and are referred to as intrinsically disordered regions (IDRs)^1,2^. They exist in a dynamic ensemble of interconverting conformational states which enables them to interact with a range of binding partners^3^ or serve as a scaffold for the association of multiple proteins^4^. The eukaryotic translation initiation factor 4G (eIF4G) is a large adaptor protein with disordered segments that acts as a framework in coordinating the assembly of initiation factors and small ribosomal subunits to initiate the process of protein synthesis^5^. A central component in this molecular organization is the heterotrimeric “eIF4F” complex formed between eIF4G, the mRNA cap-binding protein eIF4E and the helicase enzyme eIF4A^6^. This complex formation is the rate-limiting step in the regulation of cap-dependent mRNA translation and it’s aberrant activity is associated with the progression of numerous cancer associated pathologies^7,8^. Hence, multiple approaches are being undertaken to target different molecular aspects of eIF4F biology and inhibiting protein-protein interaction (PPI) between the constituent members and is an attractive avenue for the development of a comprehensive anti-cancer therapeutics^7,8^.

The IDR region of eIF4G interacts with eIF4E via a conserved canonical “Tyr-X_4_-Leu-Ø” (X is any amino acid and Ø is any hydrophobic residue) motif^9,10^. We have previously characterized the interaction of a 12mer eIF4G^D5S^ peptide with eIF4E which revealed that the recognition motif adopted the canonical “reverse L-shaped conformation” in the bound state (Figure 1A)^11^. It has a disordered N-terminus (residue 1-5) organized in an almost orthogonal orientation to the helical C-terminus (residue 6-12). The peptide forms specific hydrophilic interactions with eIF4E which includes hydrogen-bonds between Y4: P32 and L9: W73 residue pairs along with salt-bridge interaction between R6 and E132. The side-chains of L9 and L10 constitute the hydrophobic component of binding by specifically docking and interacting with the hydrophobic residues present at the interface (Figure 1A). Residue S5 (originally D in native eIF4G) present at the junction between the disordered and helical regions in the peptide was shown to be important as a helix capping residue^11^. This structural information provides an important end-state configuration of eIF4E: eIF4G PPI for deriving peptide-based inhibitors against this system^12,13^.

**Figure 1:**
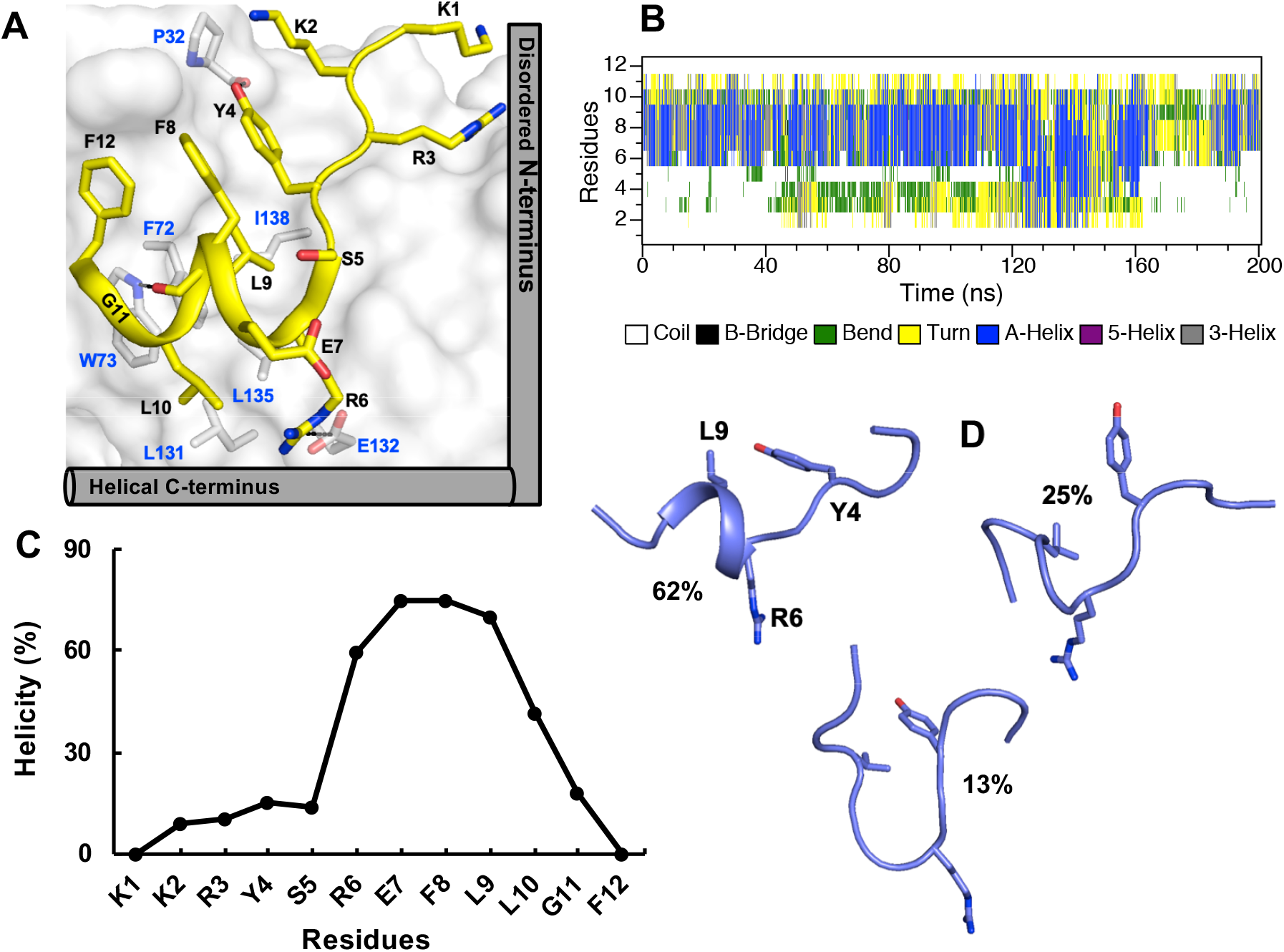
eIF4G^D5S^ peptide. **(A)** Crystal structure of 12mer eIF4G^D5S^ peptide in complex with eIF4E (PDB ID: 4BEA). The peptide backbone is shown in cartoon and side-chain in stick representation. The protein is depicted in surface and the residues involved in inter-molecular interactions with the peptide are shown as sticks. Hydrogen-bond interactions between Y4:P32 and L9:W73 along with the R6:E132 salt-bridge interaction are explicitly indicated with dashed lines. The “reverse L-shaped” conformation of the bound peptide is highlighted. **(B)** Secondary structure evolution of eIF4G^D5S^ peptide as a function of the simulation time analysed using the DSSP program^39^. **(C)** Percentage helicity of eIF4G^D5S^ peptide residues computed using “secstruct” command from the ptraj module of AMBER 18. The reported helicity is the summation of “3_10_ helix” and “α-helix” values of the individual residues. **(D)** Representative structures from three different clusters of the solution state conformations of eIF4G^D5S^ peptide. The percentage of structures in each cluster is indicated. The main-chain heavy atoms of the peptide was used for clustering the structures and it was performed using average-linkage algorithm^40^ with pairwise RMSD as a distance matrix. All the molecular graphics figures were created using PyMOL molecular visualization software (Schrödinger).

The conformational diversity of IDRs is favourable for its regulatory role, but high flexibility is not desirable in medicinal chemistry applications as only a subset of the peptide conformations will be biologically active in efficiently recognizing the specific target. Tailoring of peptide sequences for enhanced structural stability can be achieved by incorporation of conformationally-restricted non-natural amino acid derivatives^14^. In particular, substitution at the α-carbon position of the amino acid backbone is one of the most interesting and promising approaches to develop such surrogate building blocks^15^. These chemically modified residues have distinct stereochemically allowed backbone conformations with preference for specific secondary structure configurations^16,17^. α-aminoisobutyric acid (Aib) is one such non-coded amino acid derivative whose backbone structure in experimentally determined structures is largely restricted to the 3_10_/α-helical geometry^16,17^. It has been shown that introduction of Aib residue into a peptide sequence can facilitate increases in its helicity^18,19^; and that it can be employed as a general strategy to design a helical backbone scaffold for targeting PPIs^20^.

Herein, we examined the rational incorporation of Aib on the conformational properties of a peptide segment derived from the intrinsically disordered region of eIF4G and its impact on the binding affinity for eIF4E. A systematic exploration and optimization scheme was undertaken through insights from atomistic simulations which led to the development of a Aib-based peptide derivative that can specifically target the PPI between eIF4G and eIF4E for inhibition.

## Results

### Conformation of eIF4G^D5S^ peptide in solution

The conformational states of eIF4G^D5S^ peptide in explicit solvent starting from an extended structure were explored using Self-guided Langevin dynamics (SGLD)^21,22^. It is an improved sampling method which dramatically accelerates the conformational searching efficiency in biomolecular simulations and has been successfully employed to study a wide range of rare biological events within accessible simulation time scales^21,22^. Secondary structure analysis of the peptide from the generated ensemble of structures showed that it sampled a largely random conformation (Figure 1B). This is in agreement with our earlier Circular Dichroism (CD) experiment on the same peptide where it was observed to be unstructured in solution^12^. However, there was an intermittent occurrence of helical structure primarily spanning residues 6-9 during the simulation (Figure 1B). This spans a turn region and so is likely too short to be detectable from CD spectra. The average helical propensity of individual residues in the peptide showed that R6, E7, F8 and L9 adopt a helical backbone geometry in more than 50% of the structures (Figure 1C). Structure-based clustering of the conformations further indicated the presence of a metastable group (~60%) that included this helical turn (Figure 1D). Collectively, these observations suggest that there is a short local structural element with helical backbone in the free state of the eIF4G^D5S^ peptide that undergoes a transition to a higher order helix upon complexation with eIF4E as observed in the crystal structure (Figure 1A).

### TIP peptides and their binding affinity for eIF4E

We next wanted to investigate if invorporation of Aib, which is considered as a helix promoting amino acid derivative^16,17^, can influence the conformations of the peptide and their subsequent binding to eIF4E. The residues in the helical segment of the bound eIF4G^D5S^ peptide that can be potentially substituted without impacting the specificity of its interaction with eIF4E are D7, F8, G11 and F12 (Figure 1A). These residues are separated by a helical turn in the peptide sequence. We designed three dual Aib substituted derivatives by incorporating the unnatural amino acid at either i, i+4 (D7: G11 and F8: F12) or i, i+3 (F8: G11) positions and measured their affinities for eIF4E using surface plasmon resonance (SPR; Table 1 and Figure S1). TIP-01 peptide was derived by substituting Aib in the solvent exposed D7 and G11 residue positions. The binding affinity of the peptide showed about a fold improvement over the parent peptide (Table 1 and Figure S1A). The second peptide derivative “TIP-02” had Aib incorporated at residue positions F8 and F12. Interestingly, this peptide displayed an almost 80-fold decrease in binding affinity (Table 1 and Figure S1B) for eIF4E. The TIP-03 derivative with Aib substituted at residue positions F8 and G11 was an intermediate design between the previous two peptides. It had a significant improvement in binding affinity over TIP-02 but it was still about a fold lower than the parent peptide (Table 1 and Figure S1C). TIP-01 was therefore the only peptide that had a relatively better binding affinity compared to eIF4G^D5S^ whereas TIP-02 and 03 design were detrimental to eIF4E binding to different degrees.

**Table 1:**
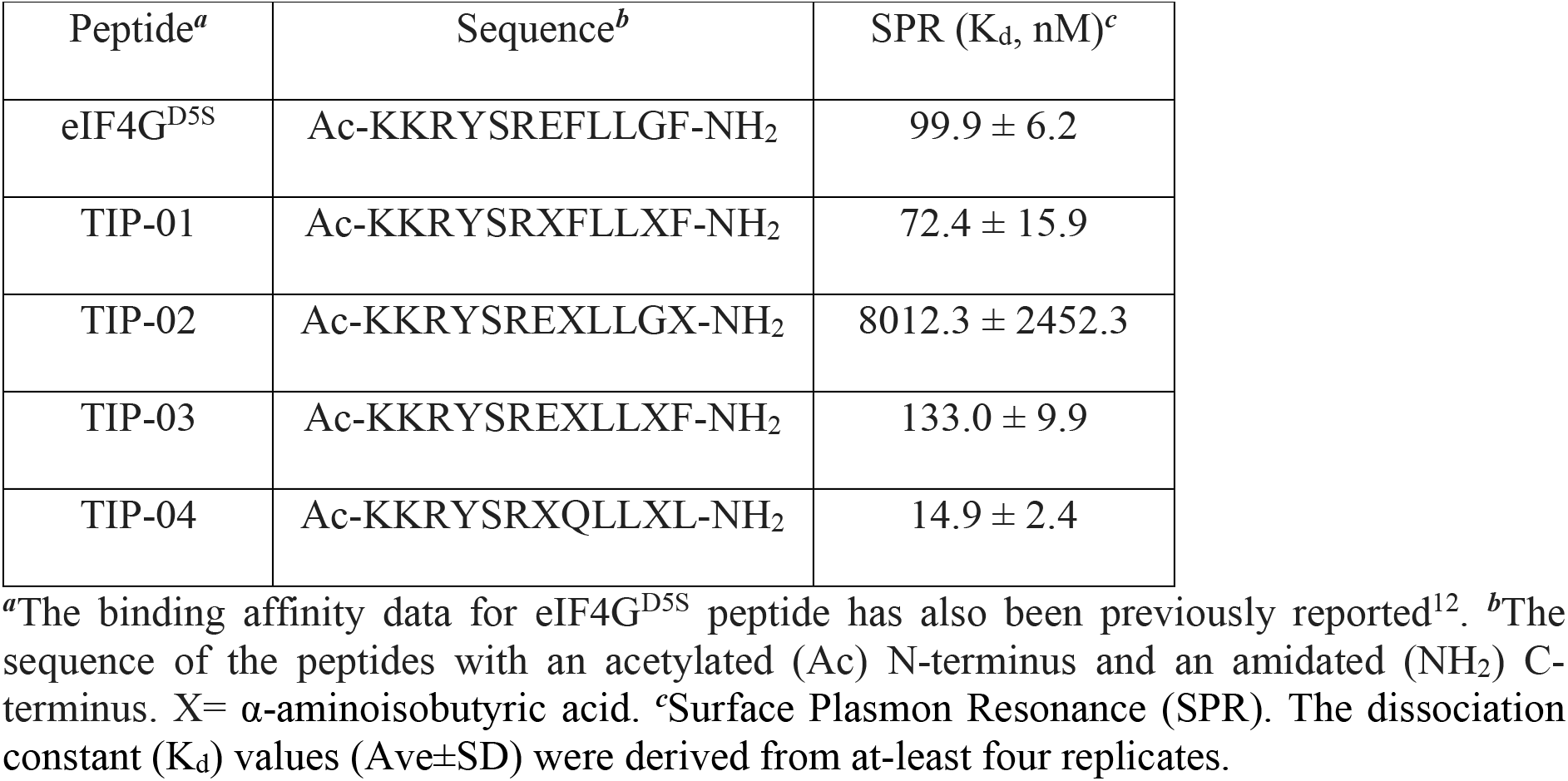
Peptides and their binding affinities^*a*^.

### Conformations of TIP peptides in solution

The secondary structure property of the three TIP peptides in solution was investigated using SGLD based sampling ^21,22^ starting from an extended state as in the case of the parent eIF4G^D5S^ peptide. TIP-01 sampled random conformations during the initial stages of the simulation but then evolved to exhibit a helical epitope from residues E7-L10 towards the C-terminal (Figures 2A and 2B). The ensemble of peptide structures were clustered into distinct groups based on the presence (~60%) or absence (~40%) of the helical turn (Figure 3A). TIP-02 unlike TIP-01, sampled random state conformation during the entire simulation (Figure 2C). There were helical signatures in the trajectory but they were not significant enough to produce at-least a 3-residue stretch with helical propensity above 50% (Figure 2D). The intrinsically disordered nature of the peptide resulted in uniformly populated (~30%) clusters in solution (Figure 3B). TIP-03 also sampled predominantly random conformations interspaced with transient helical states (Figure 2E) localized in a short three residue stretch (F8-L10; Figure 2F). The diversity in its structural configurations is also highlighted from the lack of a prevalently populated cluster (Figure 3C). Thus, this analysis showed that the Aib substitution had a relative improvement in the helical stability for TIP-01, but resulted in increased disorder in TIP-02 and TIP-03.

**Figure 2:**
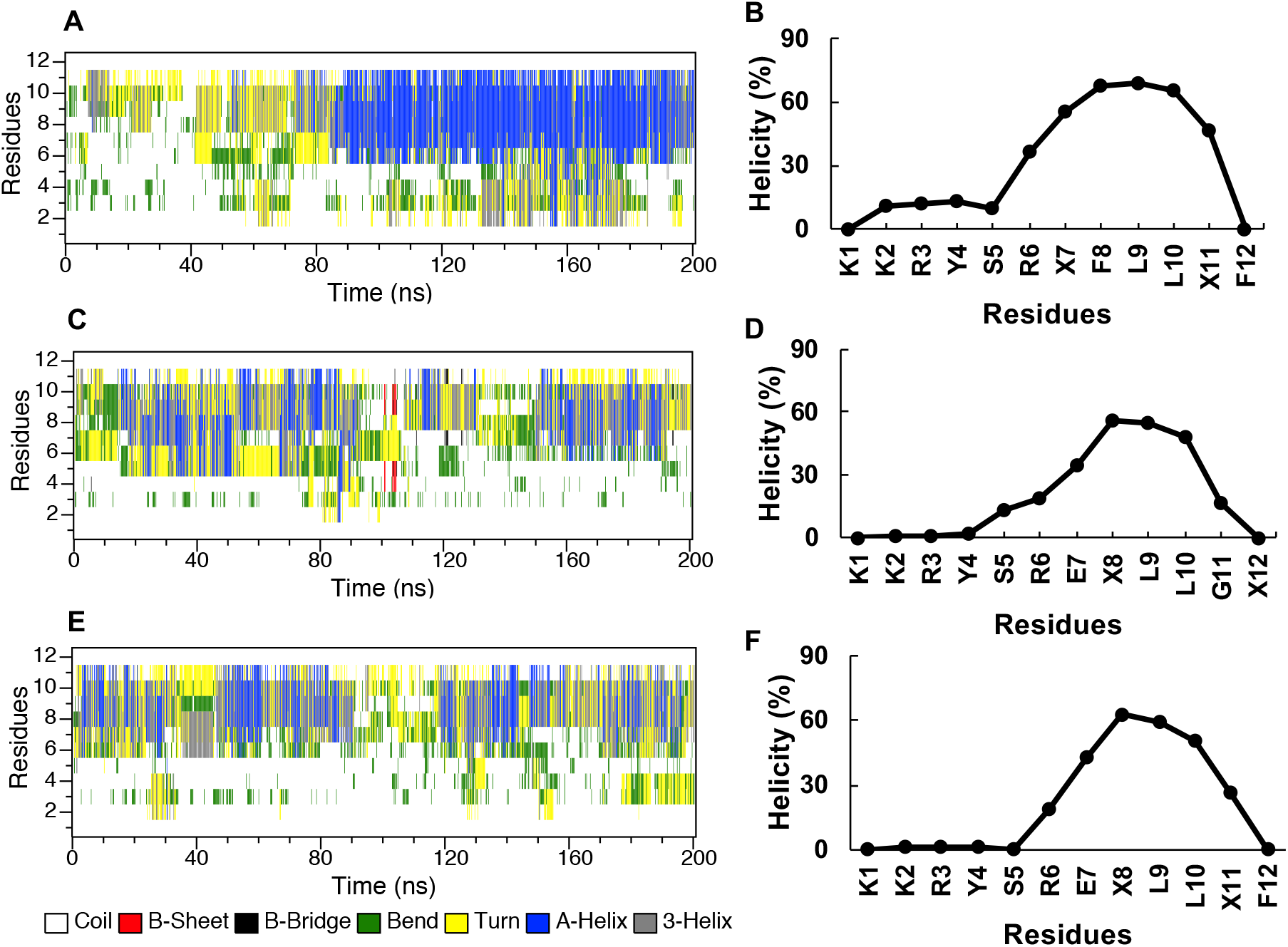
Secondary structural property of Aib peptide derivatives. Time course secondary structure evolution and percentage helicity of **(A, B)** TIP-01, **(C, D)** TIP-02 and **(D, E)** TIP-03 peptides in solution.

**Figure 3:**
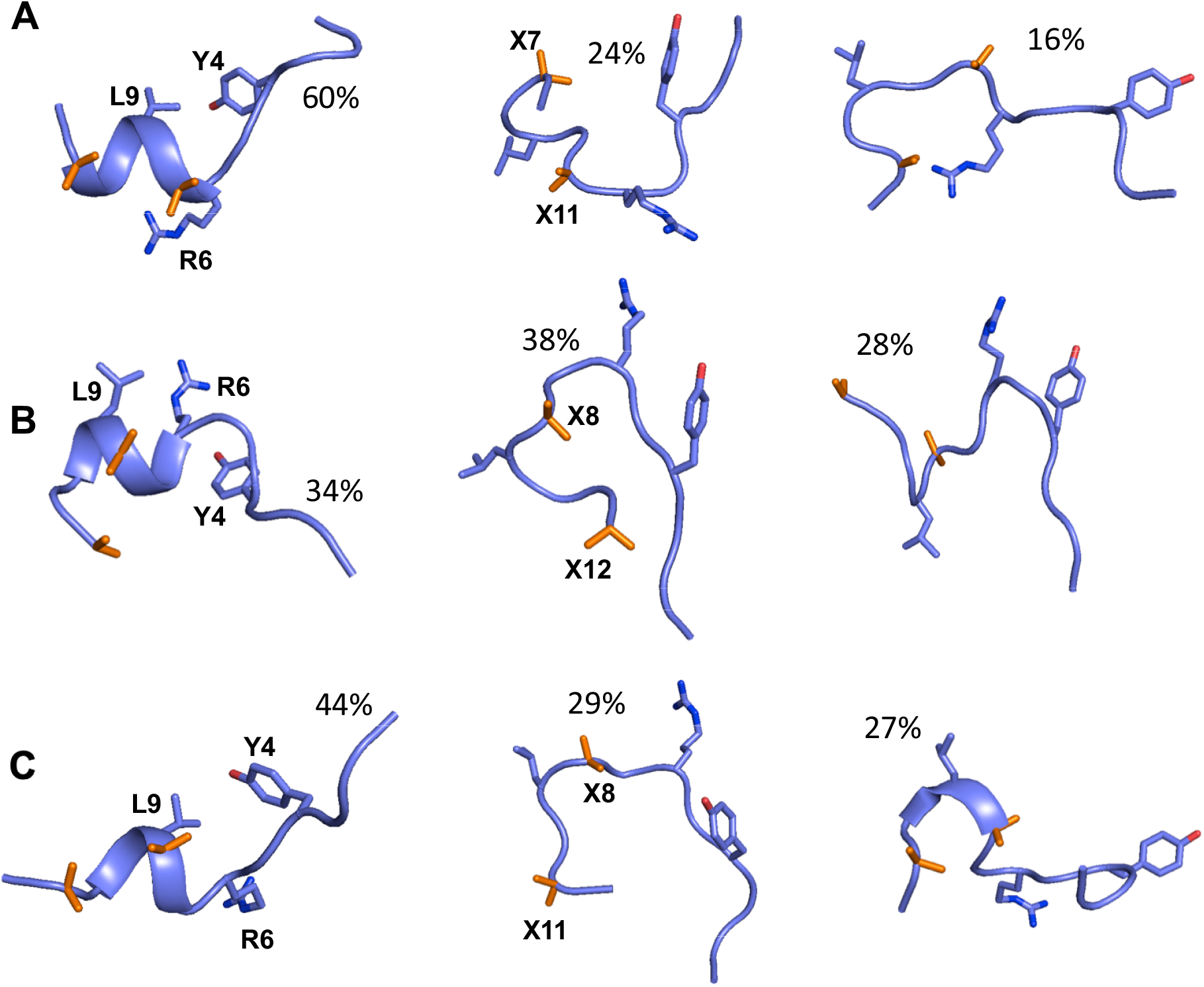
Solution state conformations of Aib peptide derivatives. Representative structures from three different cluster groups generated by clustering the ensemble of structures obtained from MD simulations of (A) TIP-01, (B) TIP-02 and (C) TIP-03 peptides in solution. The percentage of structures in each cluster is indicated. Selective residues from the peptide along with the Aib substitutions (designated by X and coloured in orange) are shown in stick representations.

### TIP peptides in the bound state with eIF4E

We also investigated the influence of Aib substitution on the eIF4E bound conformations of the peptides and the corresponding binding energetics. The complex state models of TIP peptides were generated based on the eIF4G^D5S^: eIF4E crystal structure and subjected to long-time scale (1 ms each) conventional molecular dynamics (MD) simulations. The peptides remained bound in the “Reverse L-shaped conformation” (Figure S2) and the helical segment was stable during the simulation period except for the C-terminus end of eIF4G^D5S^ and TIP-02 (Figures 4A-4D). In both these peptides, residues L10 and G11 had a reduced helical propensity (Figure 4F), which could likely have arisen from the flexibility of glycine as it is substituted by Aib in all the other peptides. Residue-wise decomposition analysis from the simulation trajectories showed that residues K1, K2, R3, Y4, R6, and L9 made significant binding energy contributions across all the peptides (Figure 4G). However, the residues from the N-terminus (K1, K2 and R3) also had a relatively higher degree of fluctuations which suggested that the interactions formed by these residues are dynamic and non-specific. Residues Y4, R6 and L9 on the other hand formed stable interactions as they constitute the specific hydrophilic and hydrophobic components of the intermolecular recognition required to interact with eIF4E (Figures S1A-S1D). E7 and G11 are solvent exposed and hence do not make any significant binding energy contributions. The three hydrophobic residues F8, L10 and F12 are all partially exposed to the solvent but they showed different energetic profiles (Figure 4G). F8 did not exhibit any significant binding energy, L10 had a reasonable contribution (~ −1 kcal/mol) while F12 displayed the most favourable (~ −2 kcal/mol) energetic contribution. In TIP-02, F12 was substituted by the shorter side-chain of Aib which resulted in the loss of the favourable hydrophobic interactions (Figure 4G). The Aib substitutions in any of the peptides do not directly contribute to the binding energy with eIF4E.

**Figure 4:**
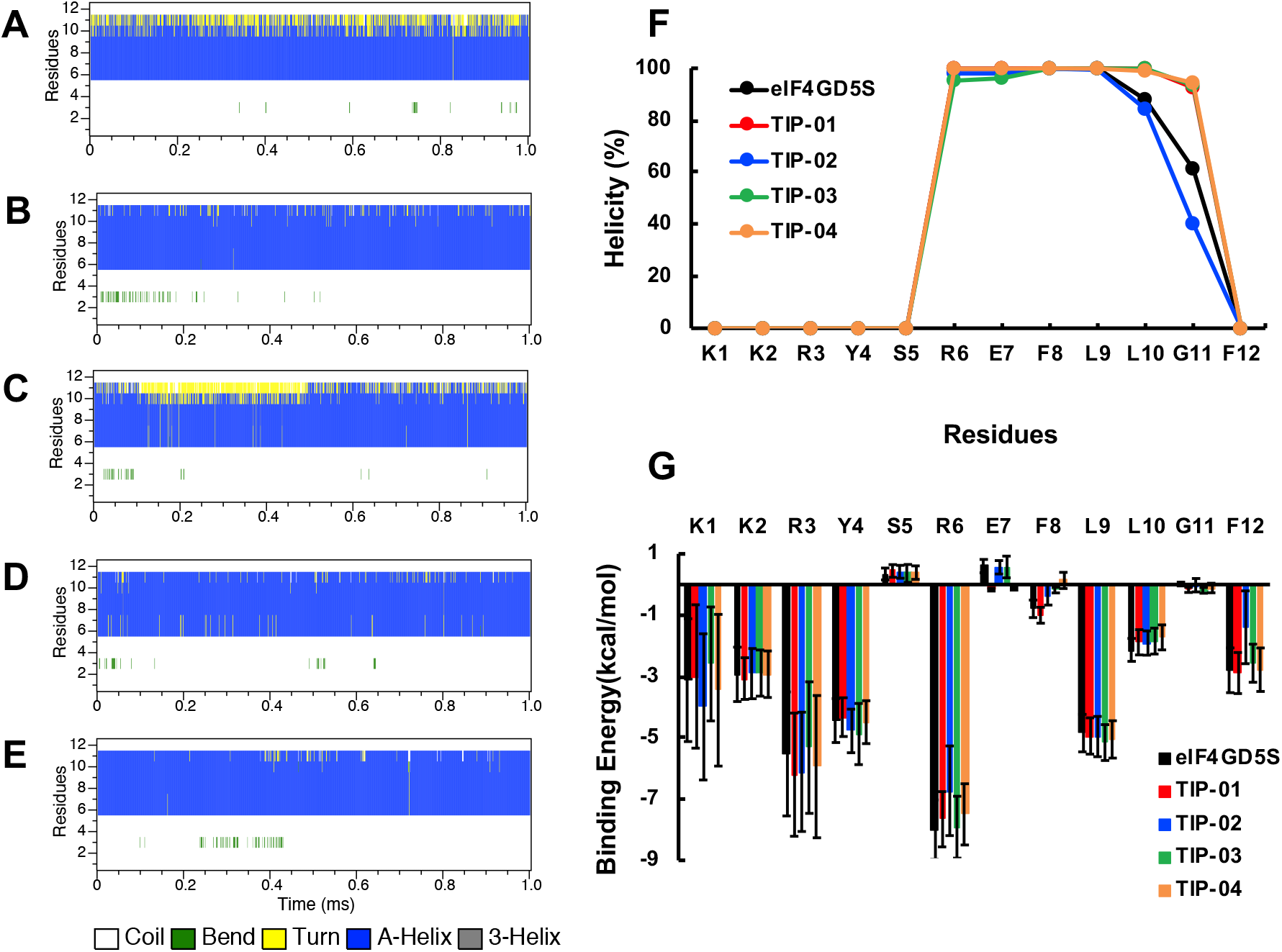
Bound state properties of the peptides. Secondary structure evolution of **(A)** eIF4G^D5S^, **(B)** TIP-01, **(C)** TIP-02, **(D)** TIP-03 and **(E)**TIP-04 peptides as a function of the simulation time. **(F)** Percentage helicity of individual residues from the different peptides. **(G)** Residue-wise binding energy contribution of the peptide in complex with eIF4E. The average binding energy and standard deviation is computed from the ensemble of structures generated from MD simulations. The calculation was done using Molecular Mechanics/Generalized Born Surface Area (MM/GBSA) method by following the same procedure and parameters as described previously^12^. The residues in the plots (**F and G**) are marked with respect to eIF4G^D5S^ peptide. See Table 1 for the Aib substituted positions in the corresponding derivatives.

### Structural reorganization of the peptides to adopt the bound state conformation

We next performed a comparative analysis between the ensemble of conformations generated from simulations of the free and bound states of the peptides by measuring their pairwise structural deviations. This variable should indicate the degree of reorganization required for the free peptides to achieve the reverse L-shaped bound conformation. The intrastate deviations of the different free peptides ranged from 0-5 Å albeit with different distribution profiles; in contrast, both the scale (0-2Å) and the nature of the distribution remained similar for the bound states across the peptides (Figures 5A-5D). Interesting data emerged with interstate comparisons where a distinct property was observed for each peptide. In eIF4G^D5S^, ~ 30% of the sub-population was in the range between 0-2Å, while the remaining (~70%) was concentrated between 2-4Å (Figure 5A) which illustrates that the majority of the peptide conformers have to undergo a higher degree of structural reorganization for complexation. In TIP-01, there was a significant jump in structural similarity between the two states of the peptide as almost half of the population (~50%) was in the range of 0-2Å (Figure 5B). This is likely due to the stable helical epitope present in significant conformers of the peptide in solution. The interstate structural variation in TIP-02 and TIP-03 peptides was considerable with the maximum (>80%) population distributed above 2Å (Figures 5C-5D) due to the intrinsically disordered nature of these peptides in solution. Thus, this analysis showed that TIP-01 would undergo the least reorganization necessary for efficiently recognizing eIF4E. We also explored the nature of specific intermolecular interactions involving residues Y4, R6 and L9 by docking representative peptide conformers in their free states onto eIF4E (Figure S3). These three residues were chosen as they define the optimum mode of interactions with eIF4E. There is a clear distinction between “eIF4G^D5S^ and TIP-01” peptides compared to “TIP-02 and TIP-03”. In the first sub-group (eIF4G^D5S^ and TIP-01), a significant population have a pre-formed helical turn in solution which enabled residues R6 and L9 to retain similar conformations between the two states of the peptides (Figures 5E and 5F). This indicated that the R6: E132 salt-bridge and the L9: W73 hydrogen-bond interactions could be formed instantly once these peptides dock into the binding interface on eIF4E. The hydrogen-bond interaction between Y4 and P32 can then be created subsequently to attain the final bound state. In the other subgroup (TIP-02 and 03), the peptides are predominantly in the disordered ensemble with low helical content which hinders their optimum docking and formation of specific intermolecular interactions. Further, even in states with the helical epitope present, TIP-02 and 03 would need the largest conformational reorganization to form the Y4: P32 hydrogen-bond (Figures 5G and 5H). This interaction is made through the disordered region of the peptide, so its formation could be a rate-limiting step in the complexation process.

**Figure 5:**
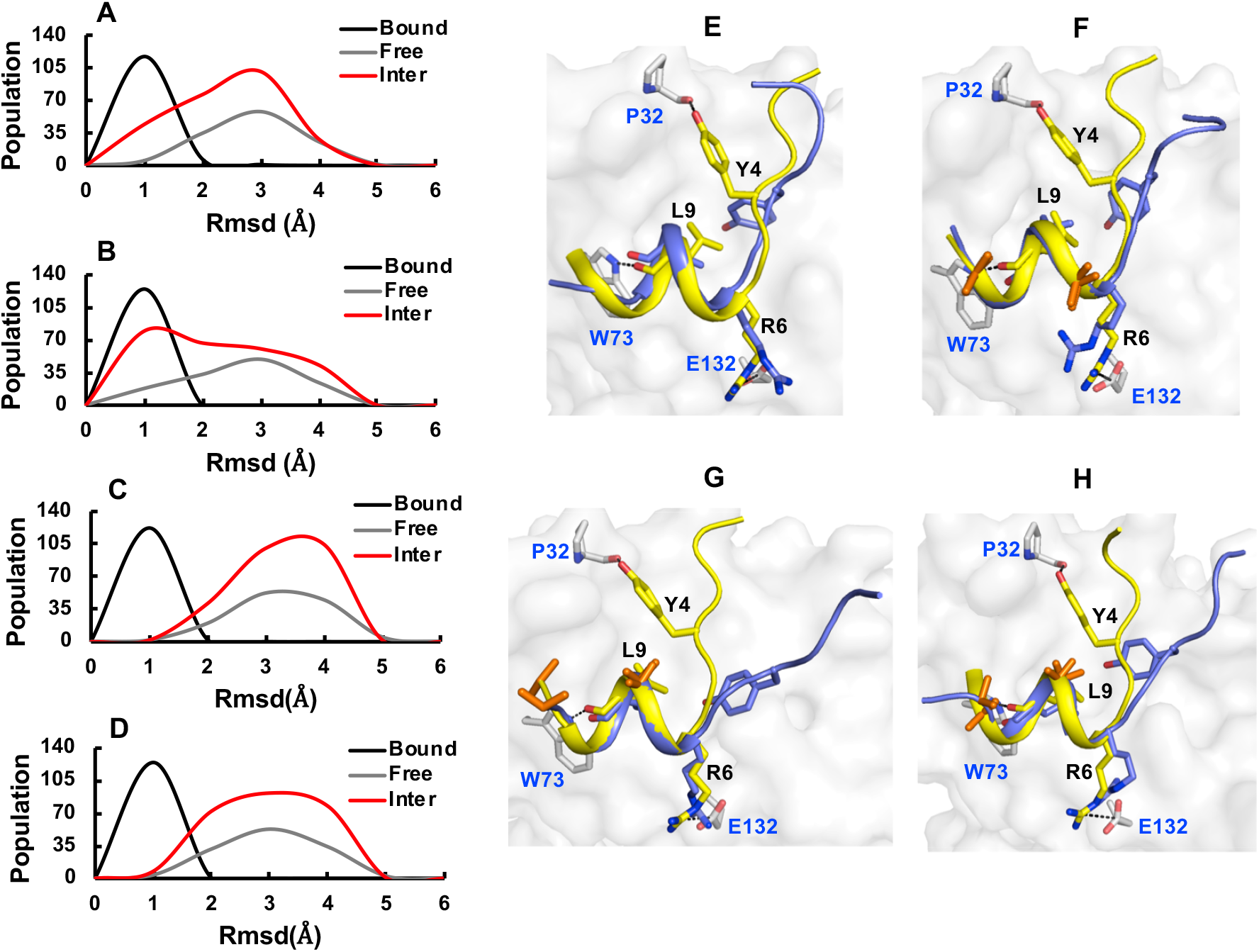
Comparison of bound and free peptides. Population distribution of **(A)** eIF4G^D5S^, **(B)** TIP-01, **(C)** TIP-02 and **(D)** TIP-03 peptide conformations in terms of the pairwise RMSD computed for the intra (bound and free) and inter (bound vs free) states of the different peptides. The main-chain heavy atoms of the peptide residues 4-12 was used for the analysis. Structural superimposition of representative bound (yellow) and free (purple) states of **(A)** eIF4G^D5S^, **(B)** TIP-01, **(C)** TIP-02 and **(D)** TIP-03 peptides. The peptides are shown in cartoon and the protein in surface representation. The side-chain of residues which are involved in specificity defining hydrogen-bond and salt-bridge interactions are explicitly depicted in stick representations. These interactions in the bound state of the peptide are highlighted with dashed lines. The side-chain of Aib substitutions are also shown as sticks coloured in orange.

### Optimization of TIP-01

TIP-01 is the only peptide that showed a relatively better binding affinity as compared to eIF4G^D5S^ (Table 1). As our analysis above suggests, the molecular basis for this could be the presence of a predominant metastable state in solution with a stable helical motif compared to the other peptides. In addition, TIP-01 also adopted a conformation which was comparatively similar to the bound state and this would lower the entropic penalty incurred upon binding. In TIP-01, hydrophilic (D7) and flexible (G11) residues were substituted by Aib. The other two peptides (TIP-02 and 03) included substitutions at F8 and F12/G11. This suggested that the two hydrophobic residue positions (F8 and F12) had an influence on the solution state conformation of the peptide. We wanted to investigate if TIP-01 can be optimized further in-order to improve its potency. Interestingly, phage display screening of eIF4E interacting peptides have previously highlighted the selective occurrence of F8Q and F12L substitutions^11^. We generated TIP-04 peptide by incorporating these substitutions into TIP-01 sequence and the peptide derivative had a ~ 7-fold improvement in binding affinity as compared to eIF4G^D5S^ (Table 1 and Figure S1D).

Conformational sampling of the peptide from an extended structure clearly indicated that it had a significantly enhanced helical character (Figure 6A). The helix, mostly across residues 6-10, was formed instantly, remained stable during the time course of the simulation (Figure 6B). MD simulation of the peptide complexed to eIF4E showed that the helicity in the bound state was also stable and the binding energetic profile of the common individual residues was similar to the other peptides (Figures 4E-4G). Neither F8Q nor F12L substitutions resulted in direct improvements in the binding energy contributions. Residues F8/Q8 does not physically interact with eIF4E, so it is energetically more favourable to incorporate a hydrophilic residue at this position. Q8 substitution also potentially stabilizes the helical conformation by reinforcing the capping role of S5 with additional hydrogen-bond interaction (Figure S4). Consequently, the bulk of the free peptide structures had pairwise deviations of <3Å, which is significantly better than TIP-01, where a sizeable population had variations above this threshold (Figures 6C and 5B). It is in-fact remarkable to observe the presence of a highly populated (~80%) peptide structure in solution which is similar to the “Reverse L-shaped conformation” observed in the bound state (Figure 6D). This suggested that the free peptide will undergo only a small amount of reorganization to efficiently interact with the protein. This is confirmed by the interstate comparison of the peptide structures where majority of the population was concentrated in the region of <2 Å (Figure 6C). In contrast, TIP-01 had a relatively widespread population distribution up to 5 Å (Figure 5B). The high degree of structural similarity in TIP-04 resulted in an effective docking posture for the disordered region in the peptide (Figure 6E) which would facilitate a faster formation of the rate-limiting hydrogen-bond interaction between Y4 and P32. Thus, in summary, TIP-04 peptide had an optimized solution state conformation which is a major molecular determinant for the improvement in the binding affinity of this peptide for eIF4E.

**Figure 6:**
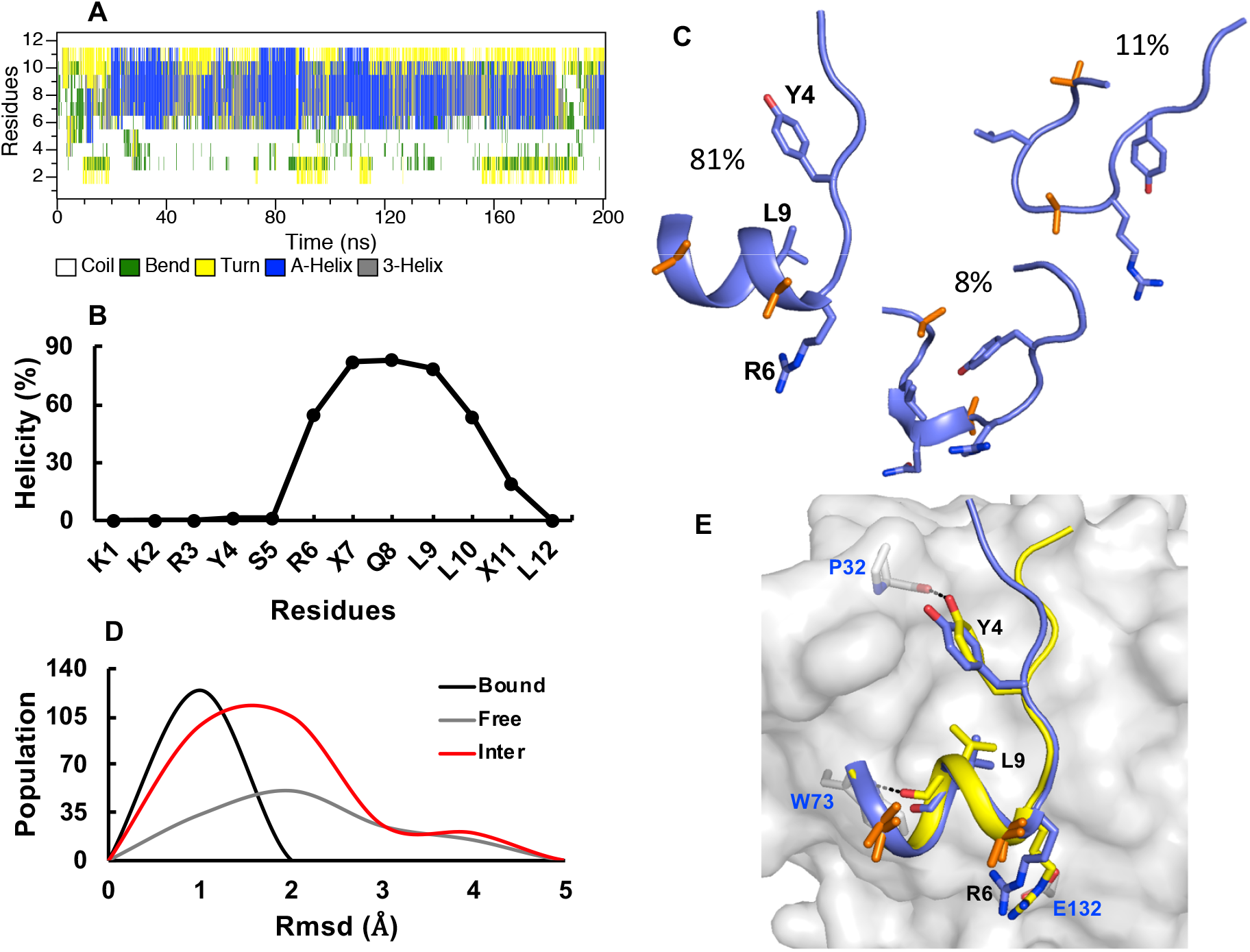
TIP-04 peptide. **(A)** Secondary structure evolution of TIP-04 peptide as a function of the simulation time. **(B)** Percentage helicity of individual residues in TIP-04 peptide. **(C)** Representative structures from three different cluster groups generated by clustering the ensemble of structures obtained from MD simulations of TIP-04 peptide. The percentage of structures in each cluster is indicated. Selective residues from the peptide along with the Aib substitutions (coloured in orange) are shown in stick representations. **(D)** Population distribution of TIP-04 peptide conformations in terms of the pairwise RMSD computed for the intra (bound and free) and inter (bound vs free) states of the peptide. **(E)** Structural superimposition of representative bound (yellow) and free (purple) states of TIP-04 peptide. The peptide is in cartoon and the protein in surface representation. The side-chain of residues which are involved in specificity defining hydrogen-bond and salt-bridge interactions are explicitly depicted in stick representations. These interactions in the bound state of the peptide are highlighted with dashed lines. The side-chain of Aib (orange) and Q8 substitutions are also shown as sticks.

## Discussion

A structure-based rational design strategy was used to develop Aib containing peptide derivatives targeted against eIF4E. The insertion of Aib into the sequence had contrasting impact on the helical property of the peptides depending on the point of incorporation. While, the helicity improved for TIP-01 and TIP-04 derivatives, it was found to be reduced in the case of TIP-02 and TIP-03. This specifies that although Aib is considered as a helix promoting non-natural amino acid^16,17^, it’s mere substitution in the sequence does not necessarily result in improved helicity. The primary structure of the parent peptide fragment derived from the IDR of eIF4G can be divided into two segments based on its chemical property; a hydrophilic N-terminus and a relatively more hydrophobic C-terminus. This difference was also manifested in its secondary structural property where the N-terminus was disordered, while localized transient helical epitopes were detected at the C-terminus. IDRs in proteins are reported to harbour such local structural elements in solution, for instance as observed in the transactivation domains of tumour suppressor p53 protein^23^. They serve as recognition motifs for interactions with binding partners like Mdm2^24^. It is interesting to note that these structural elements are selectively formed in segments within the IDR that have strategically positioned hydrophobic residues^23,24^. The incorporation of Aib residues in TIP-01 and TIP-04 systematically increases the hydrophobicity besides inducing helical backbone geometry, which collectively enhances the occurrence and stability of local helical structures in these peptides. On the contrary, Aib substitution in TIP-02 and TIP-03 reduces the overall hydrophobicity; this coupled with the presence of additional charged/flexible residues could be a primary determinant for the increase in their intrinsically disordered property in solution.

Thus, strategic incorporation of Aib along the sequence clearly is critical for the desired improvement in helicity of the peptide derivatives and their potency to interact with eIF4E. The “Reverse L-shaped conformation” is a canonical binding mode of eIF4E interacting peptides that contain the “Tyr-X4-Leu-ø” motif^25^. The specific conformation is critical as it enables the tyrosine (Tyr) residue in the motif to form a key hydrogen-bond interaction that is important for recognizing eIF4E (Figure 1A). Mutation of this residue has been shown to adversely impact the peptide’s affinity for the protein^9^. Thus, in addition to helicity, the binding activity of the peptides will also be governed by the dynamic ensemble of states that adopts bound state like conformation in solution. The mode of recognition in this system is distinct from other well studied peptide: protein systems such as p53: Mdm2^24^ or the Bcl-2 family of proteins^26^, where the critical interactions are located entirely within the helical scaffold. The conserved leucine (Leu) residue from the motif is an integral component of the helical segment across all the free peptides and it docks specifically into a pre-formed hydrophobic pocket present on the PPI interface of eIF4E (Figure 1A). A potential binding reaction would involve the formation of an initial encounter complex through selective docking of the ordered helical unit followed by structural adjustments to attain the final mode of binding. Such combined mechanism of recognition which is driven by conformational selection and optimized through an induced fit process has been shown to exist in different biomolecular interactions^27–29^. The ensemble of free TIP-04 structures are predominantly biased towards the bound state conformation which implies an efficient binding pathway for initial selection and subsequent fitting of this peptide. This will collectively decrease its entropic cost of binding and consequently as observed, TIP-04 was a potent eIF4E binder. The optimization outline of this peptide further indicates that the hinge region between the disordered and helical segment is important to preserve the bioactive conformation. In summary, TIP-04 is a viable lead compound and it’s structure-activity relationship provides important insights for consideration in the continuing efforts towards the development of peptide-based inhibitors against eIF4E in oncology.

## Methods

### Peptide synthesis

The eIF4G^D5S^ peptide, Aib substituted peptide derivatives (TIPs) and a carboxyfluorescein (FAM) labelled tracer peptide (KKRYSRDFLLALQK-(FAM)) were all ordered from and chemically synthesized by Mimotopes, Clayton, Australia. All the peptides were acetylated at the N-terminus and amidated at the C-terminus. They were purified using HPLC to more than 90% purity.

### eIF4E protein expression and purification

The full length human eIF4E protein was cloned, expressed and purified as described previously^30^.

### Surface Plasmon Resonance

Human recombinant eIF4E protein was immobilized on a CM5 sensor chip through amine coupling. The chip was first conditioned with three separate 6 s injections of 100 mM HCL, 0.1% SDS and 50 mM NaOH at a flow rate of 100 μl/min. The surface of the sensor chip was then activated with a 1:1 mixture of NHS (115 mg ml^−1^) and EDC (750 mg ml^−1^) for 7 min at 10 μl min^−1^. Purified eIF4E was diluted with 10 mM NaAc buffer (pH 5.0) to a final concentration of 0.5 μM with m^7^ GTP present in a 2:1 ratio to stabilize eIF4E. The immobilization level of eIF4E was ~1000 RU and a 7 min injection (at 10 μl min^−1^) of 1 M ethanolamine (pH 8.5) was used to block excess active coupling sites. The system was primed with an assay running buffer which consisted of 10 mM HEPES, 0.15 M NaCl, 1mM DTT and 0.1% surfactant Tween-20. The peptides were prepared by first dissolving them in 100% DMSO to a concentration of 10 mM for stock peptide solutions. The working concentration of the peptides were reached with further dilution of the stock peptide solutions into running buffer with 3% DMSO. The instrument was fully equilibrated with six buffer blanks and adding solvent correction followed by a further two blank injections. The solvent correction curve was setup by adding varying amount of 100% DMSO to 1.03x running buffer to generate a range of DMSO solutions (3.8%, 3.6%, 3.4%, 3.2%, 3%, 2.85%, 2.7% and 2.5% respectively). The peptides were injected for 60 s at a flow rate of 50 μl/min and dissociation was monitored for 180 s. SPR experiments were performed on a Biacore T100 machine and the K_d_s were calculated kinetically from the dissociation and association phase data for each peptide. The kinetic data were fitted to 1:1 binding models and each individual peptide K_d_ was determined from at-least four separate titrations.

### Modelling and simulations

The atomic coordinates of the non-natural amino acid Aib was modelled using the XLEAP module of AMBER 18^31^ and its RESP (Restrained Electrostatic Potential) based-partial charges was obtained through the R.E.D. server^32^. Other force-field parameters were derived from all-atom ff14SB^33^ force field in AMBER18. The extended conformation of the peptides with N-terminus acetylated and C-terminus amidated were generated using the TLEAP module of AMBER 18. They were solvated with TIP3P^34^ water model in a truncated octahedron box with a minimum distance of at-least 8 Å between any peptide atom and the edge of the box. The net charge of the different systems was neutralized by adding appropriate counterions. These systems were then energy minimized, heated to 300 K and equilibrated for 200 ps. The production dynamics was executed for 200 ns each for all the peptides. The generalized SGLD method (SGLDg)^21,22^ was used to sample the canonical ensemble by setting the target guiding temperature to 300 K. The local averaging time for the guiding force calculation was set to 0.2 ps, force guiding factor used to scale down low frequency energy surface was set to −0.1 and the momentum guiding factor which defines the strength of the guiding effect was set to 0.5 ps^−1^.

The crystal structure of eIF4E with eIF4G^D5S^ peptide (PDB ID: 4AZA) was used to model the bound state complexes of all the TIP peptides. The N-terminus of eIF4E was acetylated while the C-terminus was uncapped and terminated with COO^−^ carboxyl group. The terminal ends of the peptides were capped as in their free state simulations. The complexes were solvated with TIP3P water model inside a cuboid box with a minimum distance of at-least 10 Å between any solute atom and the edge of the box. The net charge of the different systems was neutralized by adding appropriate counterions. These systems were then energy minimized, heated to 300 K and equilibrated for 500 ps. Production dynamics was carried out for a period of 1μs each for all the complexes under isothermal-isobaric ensemble. The simulation temperature was maintained at 300 K through Langevin Dynamics^35^ by using a collision frequency of 1 ps^−1^ and the pressure was set to 1atm using weak-coupling^36^ with a relaxation time of 1 ps. Periodic boundary conditions were applied to all the systems and long-range electrostatic interactions were calculated using Particle Mesh Ewald (PME) method^37^. Bonds involving hydrogen atoms were constrained using SHAKE^38^ and the integration time step was set to 2 fs.

## Acknowledgments

DPL and DL are supported by VR grant 2013_08807. DL gratefully acknowledges the technical support from members of the Biocomputing Centre at Bioinformatics Institute, A*STAR Singapore especially Yong Tai Pang, Toe Chin Siang and Liang Zhu.

