## Supplementary Information for "Conformational ordering of intrinsically disordered peptides for targeting translation initiation"

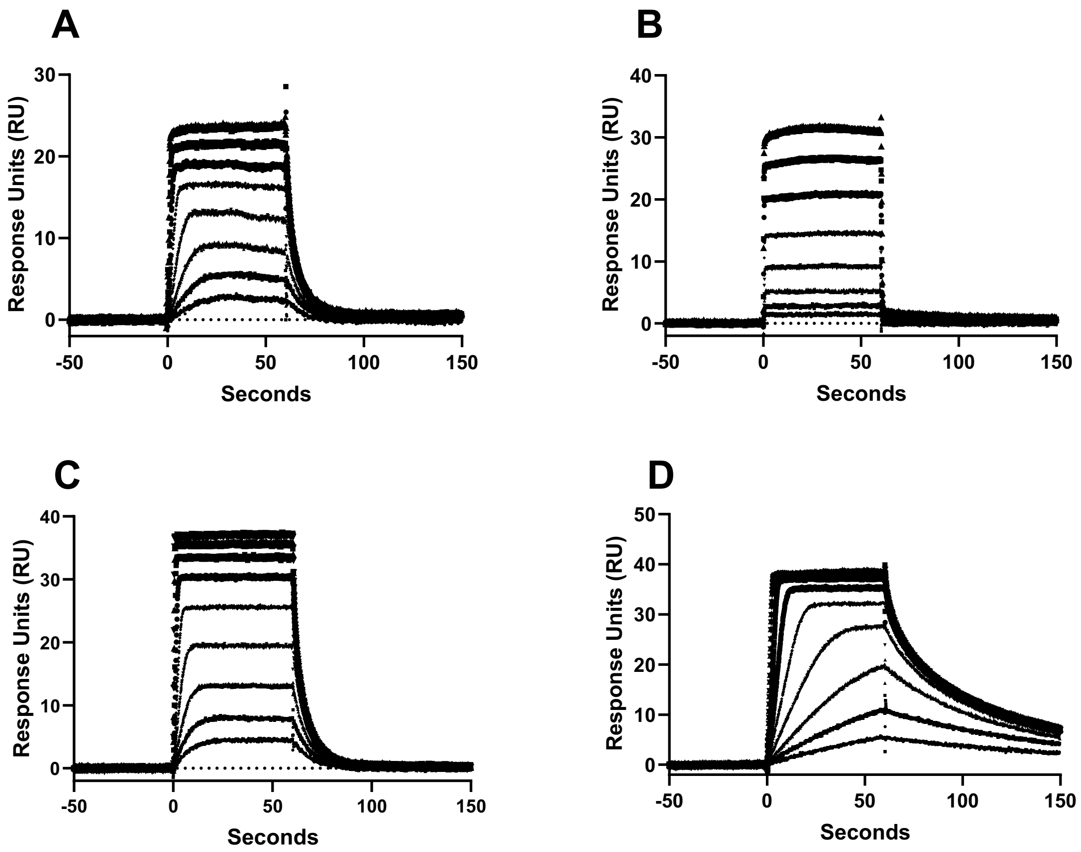


**Figure S1: Sensograms.** Representative sensogram traces for **(A)** TIP-01, **(B)** TIP-02, **(C)** TIP-03 and **(D)** TIP-04 from surface plasmon resonance (SPR) experiments.

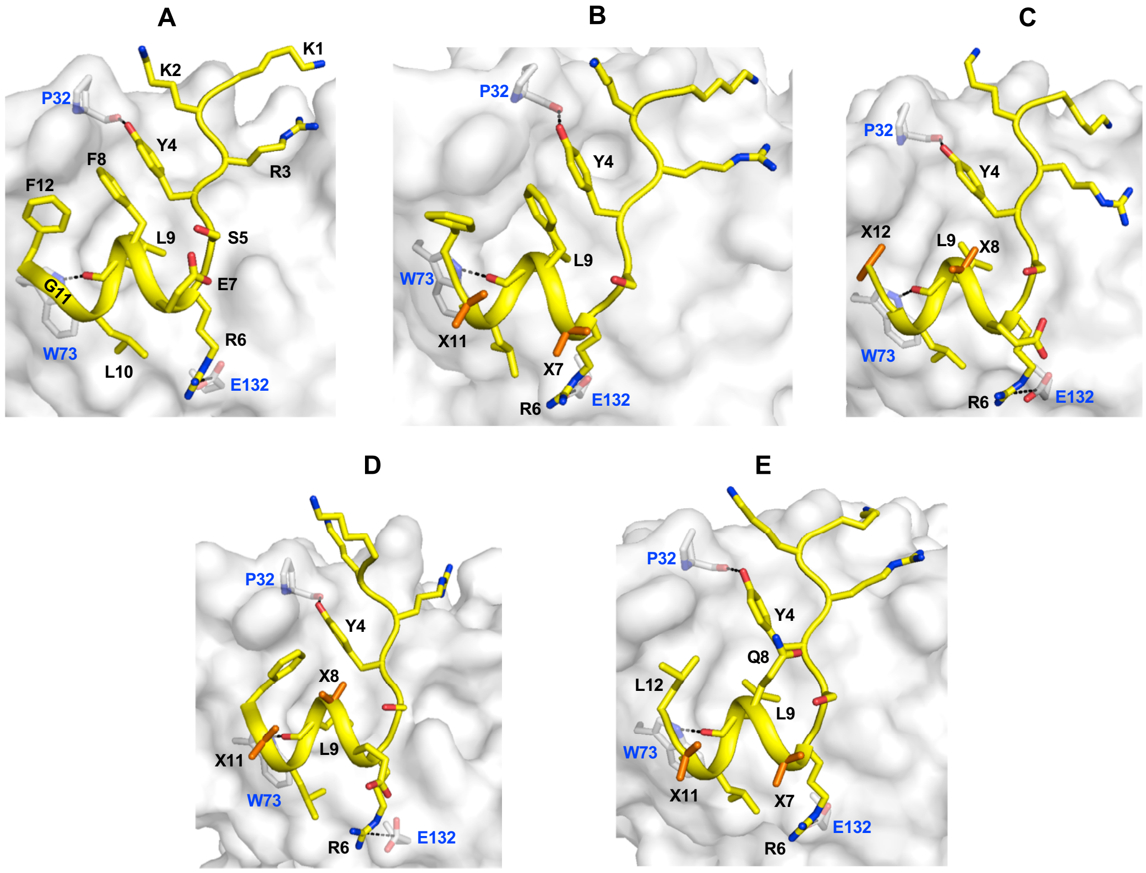


**Figure S2: Bound state of the peptides.** Representative eIF4E bound state structures of **(A)** eIF4G$^{D5S}$, **(B)** TIP-01, **(C)** TIP-02, (D) TIP-03 and **(D)** TIP-04 peptides derived from MD simulations of the respective complex. The representative structures were selected from the highest populated (> 80%) cluster for each system. The main-chain heavy atoms of the peptide were used for clustering the structures. The peptide backbone is shown in cartoon and side-chain in stick representation. The protein is depicted in surface and the residues involved in inter-molecular interactions with the peptide are shown as sticks. Hydrogen-bond interactions between Y4:P32 and L9:W73 along with the R6:E132 salt-bridge interaction are explicitly indicated with dashed lines. The side-chain of Aib substitutions (designated with symbol X) are also shown as sticks coloured in orange.

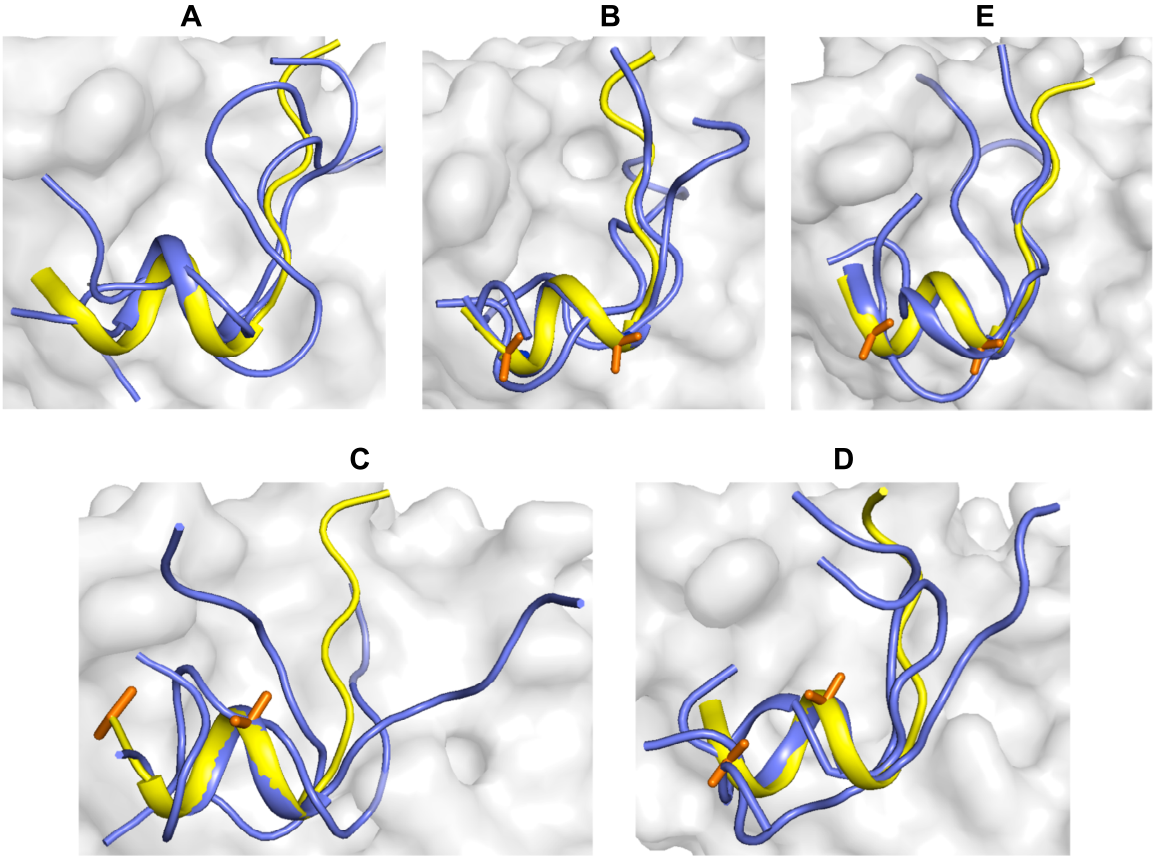


**Figure S2: Comparison of bound and free peptides.** Structural superimposition of representative bound (yellow) and free (purple) states of **(A)** eIF4G$^{D5S}$, **(B)** TIP-01, **(C)** TIP-02 and **(D)** TIP-03 and **(D)** TIP-04 peptides. The peptides are shown in cartoon and the protein in surface representation. The representative structures of the bound state is selected from the highest populated (> 80%) cluster for each system (See Figure S1). The representative structures of the peptides was selected from three different cluster groups (See Figure 1, 2 and 6). The main-chain heavy atoms of the peptide was used for clustering the structures.

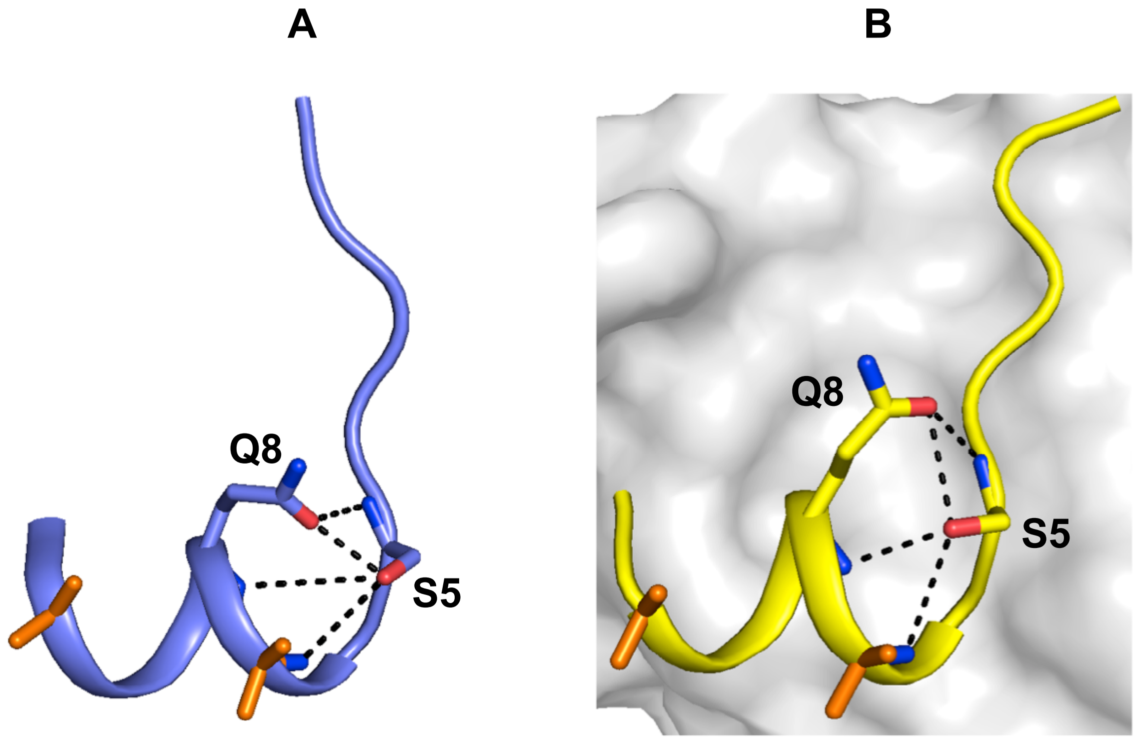


**Figure S3: F8Q substitution. (A, B)** Representative structure of TIP-04 in free and bound states respectively. The peptide is shown in cartoon and the protein in surface representation. The potential hydrogen-bond interactions formed by the side-chain of residues S5 and Q8 with the backbone atoms are indicated with dotted lines. The side-chain of Aib substituted residues are shown in stick representation coloured in orange.
